# Antibacterial wound dressing enables reliable detection of uric acid in exudates

**DOI:** 10.1101/2025.09.29.679342

**Authors:** Bozhang Li, Mingrui Lv

## Abstract

Through the successful grafting of epoxypropyl dimethyl dodecyl ammonium chloride (EDDAC) onto bacterial cellulose (BC) and multi-walled carbon nanotube (CNT) surfaces, a novel and stable antibacterial wound dressing (BC-CNTs-EDDAC) was fabricated. Compared with bare BC membranes, BC-EDDAC exhibited significantly enhanced antibacterial activity. Furthermore, a non-enzymatic uric acid sensor was constructed on the membrane surface via the self-assembly of AuNps, resulting in an antibacterial wound dressing with integrated UA-sensing capability. This design provides a new strategy for both promoting wound healing and real-time assessment of wound recovery status.

## 1. INTRODUCTION

Traditional wound dressings, represented by gauze and cotton, primarily function to cover the wound surface [1], absorb exudates, and prevent contamination [2], offering the advantages of low cost and ease of use. However, they only provide passive protection, are prone to adhering to the wound [3], and may cause secondary injury and pain during replacement [4]. Moreover, conventional dressings merely serve covering and absorption roles, lacking the capability to monitor the wound healing process in real time [5]. Physicians or patients often need to frequently remove the dressing and rely on visual inspection to judge infection [6] or healing progress, a method dependent on subjective experience and incapable of accurately assessing whether the wound is in the inflammatory, proliferative, or remodeling phase [7]. This process not only disrupts the moist healing environment but also increases the risks of secondary injury and infection [8]. Consequently, traditional dressings are insufficient for dynamically and precisely reflecting wound recovery stages [9], limiting personalized and efficient wound management, particularly in the care of chronic or complex wounds.

Uric acid (UA), a product of purine metabolism [10], is elevated in wound exudates when rapid cellular metabolism and tissue degradation occur [11]. High levels of UA can trigger local inflammatory responses, exacerbate the inflammatory microenvironment and impair wound healing [12]. Studies have revealed that changes in metabolites such as UA in wound exudates may serve as potential biomarkers for infection monitoring and delayed healing [13].

In modern healthcare, the development of advanced wound care solutions is essential to address the complexity of wound management. Integrating antibacterial properties, UA monitoring, and wound healing promotion into a single dressing not only provides protection against infection but also enables real-time monitoring of UA levels in wound exudates, offering valuable insights into the metabolic state and healing progress of the wound [14]. The convergence of antibacterial materials and wearable sensor technology heralds a new era in wound management. The development of wearable sensors capable of functioning as antibacterial wound dressings is driven by the urgent need for real-time monitoring and infection control [15].

Bacterial cellulose (BC) offers remarkable advantages as a wound dressing: its three-dimensional nanofiber network retains high water content [16], ensuring a moist environment that promotes cell migration; it also possesses excellent mechanical properties and flexibility [17], allowing it to conform to irregular wound surfaces. With high purity and outstanding biocompatibility, BC is less likely to trigger inflammatory responses [18], while its superior breathability and fluid-handling capacity further enhance its performance [19]. In addition, BC surfaces are readily functionalized [20], enabling drug loading or incorporation of antibacterial agents. Thus, BC is regarded as an ideal material for next-generation wound dressings. Epoxypropyl dimethyl dodecyl ammonium chloride (EDDAC), a quaternary ammonium antibacterial agent [21], exhibits broad-spectrum and highly efficient bactericidal activity against common pathogens such as Staphylococcus aureus and Escherichia coli. Its epoxy groups can covalently bond with BC, ensuring firm immobilization, reducing side effects from leaching, and providing long-term stable antibacterial activity, making it suitable for chronic wound or postoperative incision dressings.

Uric acid can be catalytically oxidized at a specific potential in the presence of deposited gold nanoparticles (AuNPs), producing allantoin, carbon dioxide, and ammonia [22]. When AuNPs are immobilized on BC, the three-dimensional nanofiber network of BC provides a high surface area and stable support, preventing aggregation. UA is efficiently catalyzed and oxidized on this surface, generating significant current signals and enabling sensitive and selective UA detection. Carbon nanotubes (CNTs), with their exceptional electrical conductivity [23], form efficient electron transport bridges between the BC network and AuNPs, thereby enhancing electrochemical responses, lowering detection limits, and allowing rapid, stable, and reliable detection of small molecules such as UA.

This study aims to develop an efficient biosensor and introduces a novel composite material “BC-EDDAC-CNTs-AuNPs” for monitoring UA concentrations in wound exudates while simultaneously exhibiting excellent antibacterial properties. This unique material design offers new insights into biosensor development, with the potential to improve the quality of life of patients with chronic wounds and alleviate the burden on healthcare systems.

## 2. Experimental section

### 2.1. Chemicals

BC, EDDAC, CNTs Epichlorohydrin,Chloroauric acid were procured from Sigma-Aldrich (USA). Electrochemical experiments utilized a 10 mM PBS buffer (pH 7.4) containing 100 mM potassium chloride and 10 mM ferricyanide/ferrocyanide ([Fe (CN)_6_] ^3−^ /^4−^).

### 2.2. Instruments

The experimental instruments used in this study included Fourier-transform infrared spectroscopy (FTIR, Bruker Optics, Billerica, MA), a scanning electron microscope (SEM, JSM-7500F, Hitachi, Japan), and an electrochemical workstation (CHI 660E), where a printed electrode system was employed with BC-EDDAC-CNTs-AuNPs as the working electrode, a silver electrode as the counter electrode, and a carbon paste electrode as the reference electrode. In addition, a high-speed disperser (T25 DS25, IKA, Germany) was utilized during sample preparation.

### 2.3. Preparation of the BC-EDDAC film

EDDAC was prepared through the reaction of dimethyldodecylamine (0.01 mmol) with epichlorohydrin (0.04 mmol) at 80 °C for 1 h. The reaction mixture gradually changed in color from clear to dark yellow, indicating product formation. The resulting product was precipitated using diethyl ether, and the crude EDDAC was isolated via centrifugation. To purify the material, the precipitate was washed three times with diethyl ether under ultrasonic agitation, followed by vacuum drying. For surface functionalization, BC membranes were immersed in a 0.06 mol/L aqueous EDDAC solution and allowed to react at ambient temperature (Figure 1). Finally, the membranes were thoroughly rinsed with deionized water.

**Figure 1.**
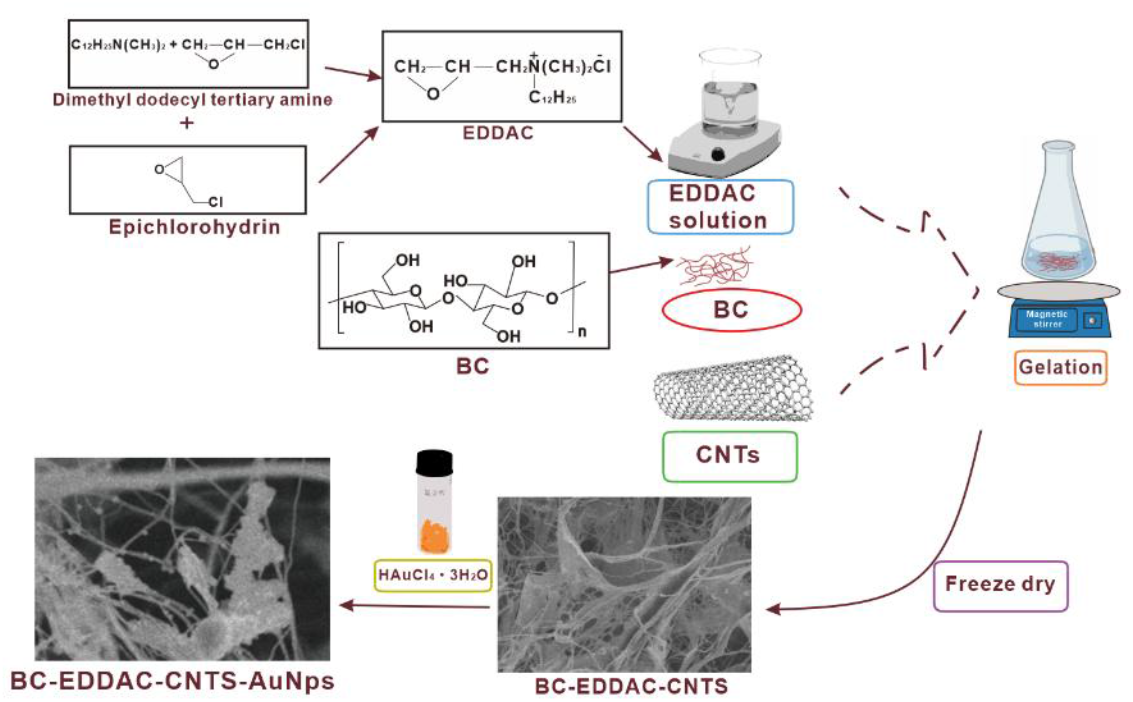
Synthesis process of BC-EDDAC-CNTS-AuNps.

### 2.4. Preparation of the BC-EDDAC -CNTs solution

Acid-treated CNTs were first pre-dispersed in distilled water using an ultrasonic generator. The BC-EDDAC film was cut into small fragments and further disintegrated with a homogenizer. Excess water in the BC-EDDAC fragments was gently removed, after which the fragments were immersed in a mixed solution containing NaOH, urea, distilled water, and CNTs. The suspension was pre-cooled at −6 °C for 2 h, followed by stirring at ambient temperature for approximately 5 min.

### 2.5. Gelation of the BC-EDDAC-CNTS Solution

Epichlorohydrin was employed as a crosslinking agent for the gel formation of the BC-EDDAC-CNTs solution. A defined amount of epichlorohydrin was slowly added dropwise into the solution under vigorous stirring at 2,500 rpm. The resulting mixture was then transferred to an oven and maintained at 60 °C overnight to allow complete crosslinking. After the reaction, the product was thoroughly washed with excess distilled water to remove residual reagents, and subsequently freeze-dried to obtain the BC-EDDAC-CNTs composite film.

### 2.6. Fabrication of BC-EDDAC-CNTS-AuNps film

AuNPs were electrodeposited onto the BC-EDDAC-CNTs surface by drop-casting 40 µL of 3.0 mM HAuCl_4_·3H_2_O solution, followed by applying a potential of −0.6 V (compare to Ag reference electrode) for 120 s. Prior to use, the AuNPs-modified composite films were thoroughly rinsed with deionized water and subsequently dried at 60 °C for 1 h.

## 3. Results and discussions

### 3.1. Characterization of the BC-EDDAC-CNTS-AuNps

Figure 2a and 2a_1_ present the SEM morphology of freeze-dried BC and BC-EDDAC hydrogels, where BC exhibits a well-defined three-dimensional nanofibrous network. After the incorporation of CNTs, the SEM images in Figure 2b and 2b_1_ clearly demonstrate that CNTs were successfully embedded within the BC network. When BC-CNTs were physically blended with 0.3 wt% EDDAC, the resulting BC-CNTs-EDDAC hydrogel retained the interconnected three-dimensional porous structure (Figure 2c and 2c_1_). The presence of hydrophilic groups in EDDAC, along with its incorporation into the BC-CNTs network, further enhanced the hydrophilicity of the hydrogel. Such a porous and interconnected architecture is favorable for nutrient and oxygen diffusion, thereby supporting wound healing. Moreover, Figure 2d and 2d_1_ provide clear evidence that AuNPs were successfully deposited on the surface of the BC-CNTs-EDDAC hydrogel.

**Figure 2.**
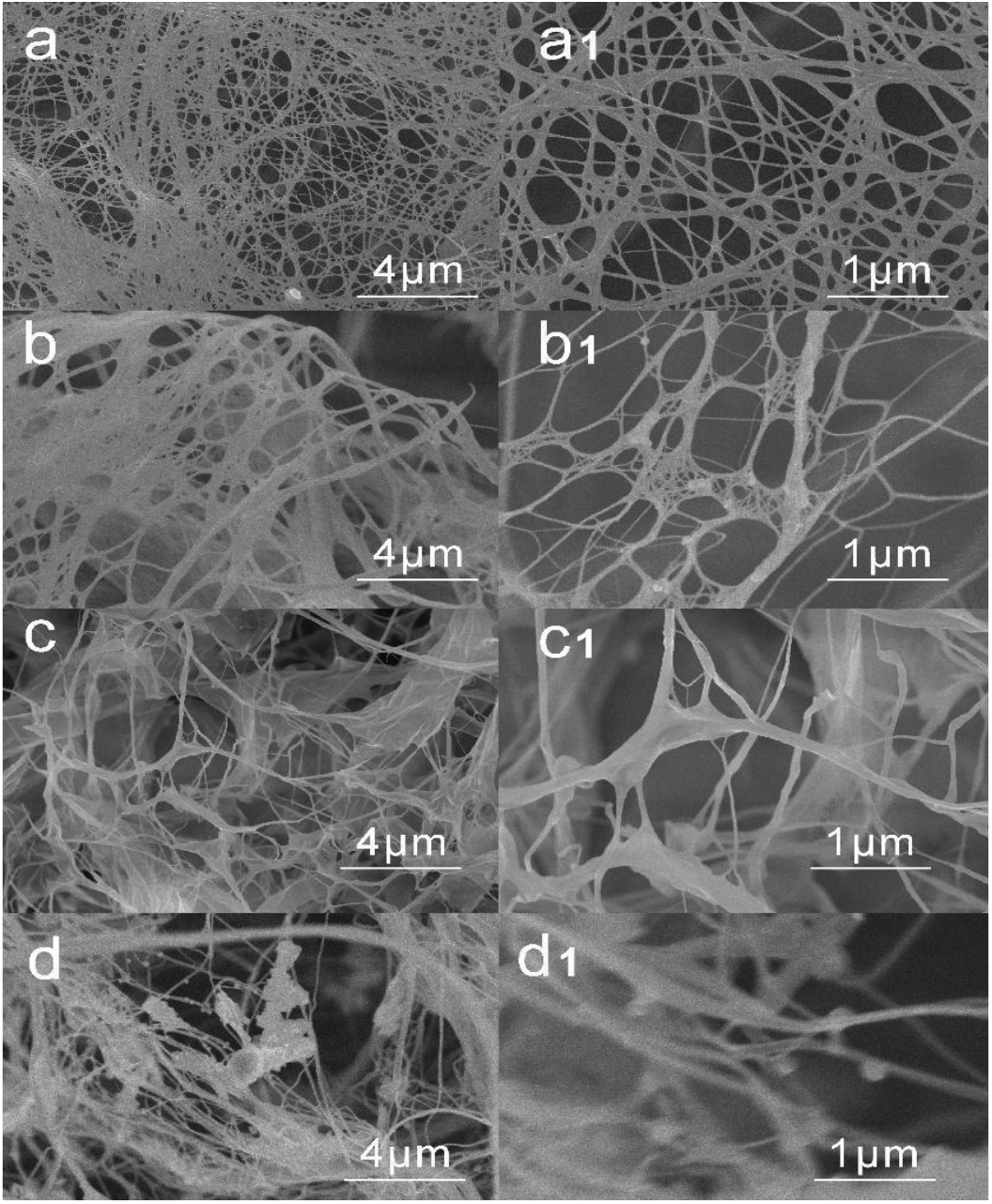
SEM images of (a) BC, (a_1_) the magnified figure of BC, (b) BC-CNTS, (b_1_) the magnified figure of BC-CNTS, (c) BC-EDDAC-CNTS, (c_1_) the magnified figure of BC-EDDAC-CNTS, (d) BC-EDDAC-CNTS-AuNps, (d_1_) the magnified figure of BC-EDDAC-CNTS-Au Nps.

**Figure 3.**
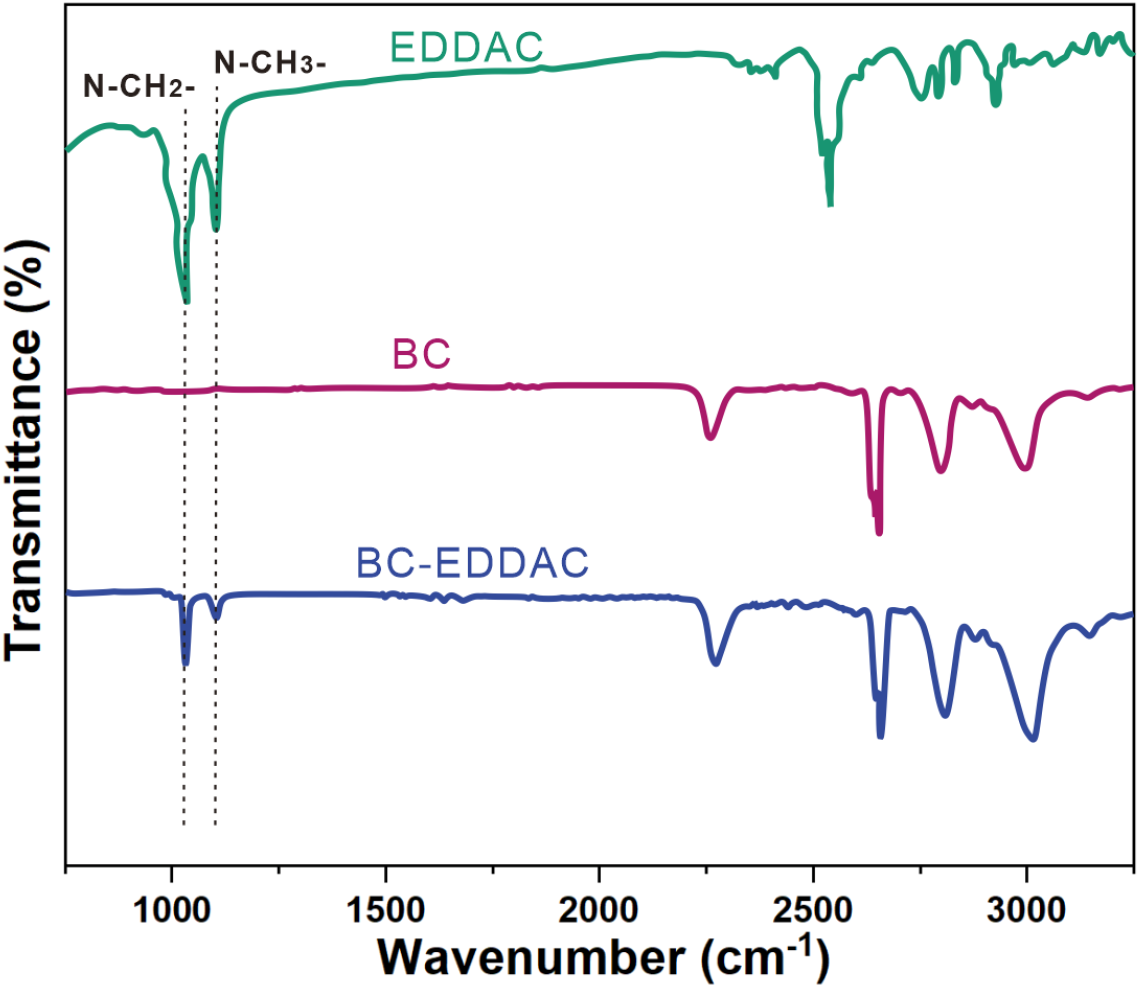
FTIR spectra of BC, EDDAC, BC-EDDAC.

FTIR was employed to characterize the BC and BC-EDDAC membranes in order to investigate structural changes following the incorporation of quaternary ammonium groups. The spectra of BC-EDDAC exhibited characteristic absorption peaks corresponding to both BC and EDDAC, with slight shifts observed in some peak positions. These results confirm that EDDAC was successfully incorporated into the BC matrix, leading to the formation of the BC-EDDAC composite.

### 3.2. Antibacterial property of BC-EDDAC

Since the amount of EDDAC plays a critical role in determining the antibacterial performance of BC-EDDAC, we evaluated the ratio of BC to EDDAC. In this part of the study, the antibacterial activity of BC and BC-EDDAC-CNTs composites against E. coli and S. aureus was quantitatively assessed using the plate colony counting method after 3 h of incubation. The specific results are presented below.

The antibacterial effects of BC and BC-EDDAC composite materials against E. coli and S. aureus over a 3-hour period are shown in figure. 4. It is evident that there are more bacteria in the BC composite material, whereas the BC-EDDAC composite materials have fewer bacteria, especially in BC-0.3 EDDAC, where almost no bacteria are present. The antibacterial results show that BC-EDDAC exhibits an inhibition rate of over 99% against both E. coli and S. aureus, demonstrating excellent antibacterial performance. In the subsequent 24-hour sampling, bacteria in the BC composite material have multiplied significantly, while the BC-EDDAC composite material shows almost no colony formation, indicating that bacteria have become largely deactivated after 3 hours of exposure. This is attributed to the fact that the isoelectric point (pH) of bacteria is generally around 2-5, and the bacterial cell membrane surface contains numerous acidic and strong phosphate groups, making the bacterial surface negatively charged. On the other hand, EDDAC molecules themselves contain positively charged amino groups. Therefore, when EDDAC comes into contact with bacteria, EDDAC readily adsorbs to the bacteria, damaging the bacterial cell wall and causing the release of components such as proteins, leading to bacterial death. Additionally, EDDAC can enter the interior of the bacteria, cause protein aggregation and disrupting the normal reproductive environment of the bacteria, ultimately leading to bacterial demise.

**Figure 4.**
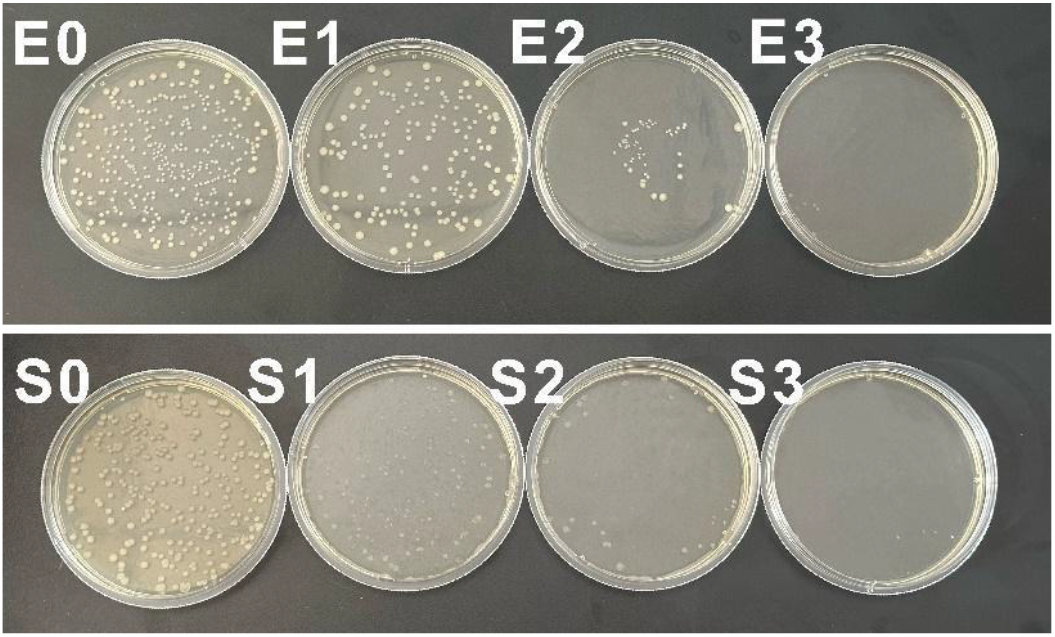
3 h antibacterial effect photo of BC (E1), BC-0.1 EDDAC (E2) and BC-0.3 EDDAC (E3) against E. coli (up) and S.aureus (down).

### 3.3. Detect of UA

The CV and DPV curves (Figure 5a, 5b) reveal a distinct oxidation response to UA compared with PBS, confirming the sensor’s strong sensitivity. Moreover, as shown in Figure 5c, the presence of common interfering species such as AA, LA, and Glu does not significantly affect the UA signal, indicating that the sensor maintains a clear and stable response to UA. These results demonstrate the excellent selectivity and anti-interference capability of the BC-EDDAC-based sensor for UA detection in complex environments.

**Figure 5.**
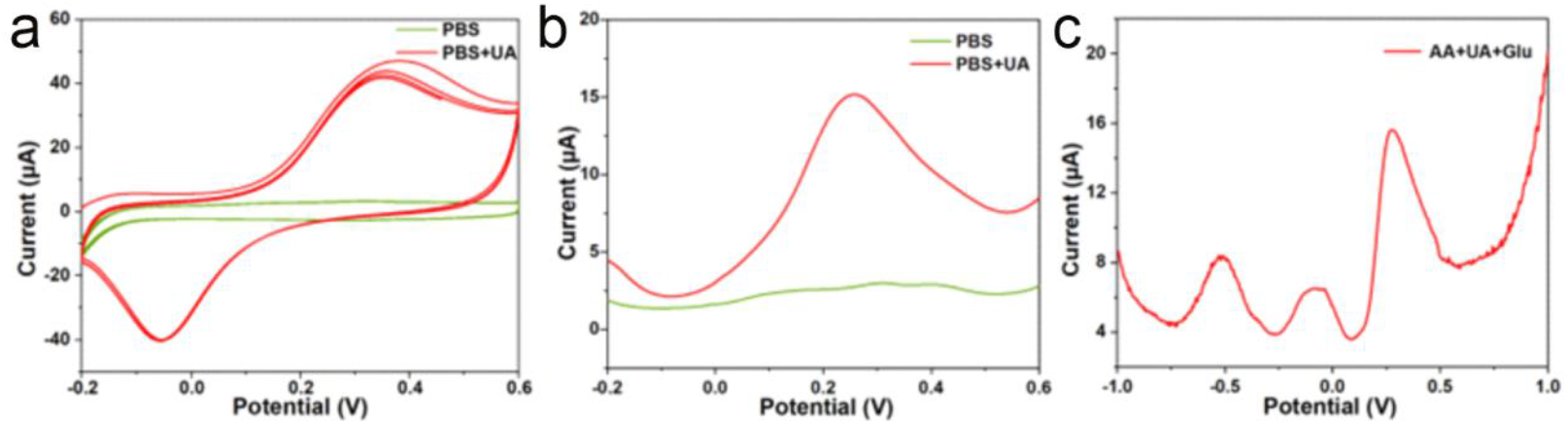
(a) CV and (b) DPV profiles of BC-EDDAC in PBS and in PBS containing 100 μM UA. (c) Selective response of the sensing layer toward UA (10 μM), AA (100 μM), and LA (100 μM) in PBS at an applied potential of 0.28 V.

The I–T responses (Figure 6a) show stepwise current increases with successive UA additions, demonstrating sensitive detection. The corresponding calibration plot (Figure 6b) reveals a strong linear relationship between current response and UA concentration, with regression equation ΔI = 0.07C_UA_ + 9.00 and an excellent correlation coefficient (R^2^ = 0.995). The inset further confirms reliable linearity in the low concentration range (R^2^ = 0.992). These results indicate that the sensor exhibits outstanding linearity and accuracy for UA quantification across a broad concentration range.

**Figure 6.**
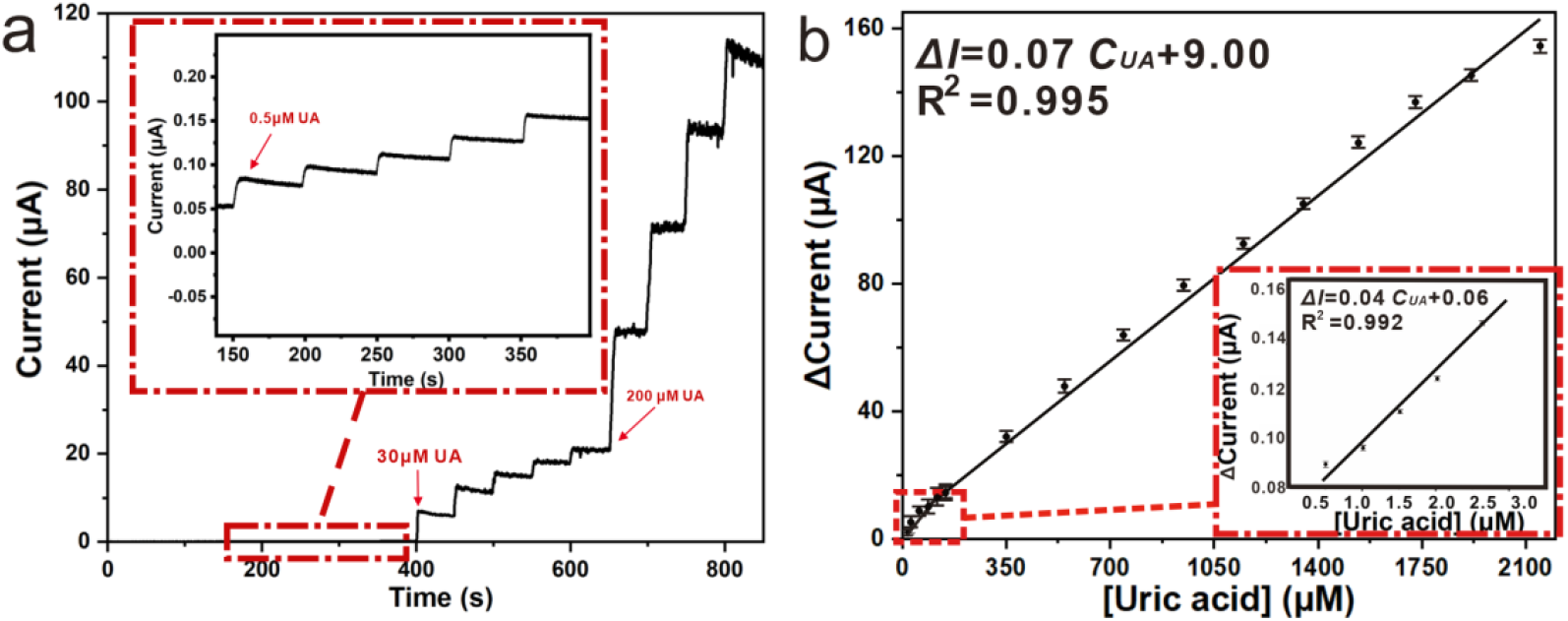
(a) Sensor I–T responses to incremental UA additions in PBS at 0.28 V, and (b) the resulting calibration plot.

## 4. Conclusion

In conclusion, we developed a BC-based wound dressing capable of detecting uric acid in wound exudates. The synergistic combination of BC and EDDAC endowed the material with remarkable antibacterial activity, while the incorporation of AuNPs enabled the biosensor to achieve high precision in UA detection. This feature is essential for tracking and assessing critical phases of wound repair, such as epidermal proliferation, aberrant differentiation, angiogenesis, immune activity, and inflammatory processes. The fabricated BC-EDDAC dressing, equipped with advanced biosensing capability, is designed to achieve optimal wound management and ensure stable and reliable performance across diverse clinical settings.

